# From Field to FASTA: Onsite DNA Barcoding the Mushrooms of the 2024 NAMA Foray

**DOI:** 10.1101/2025.07.16.664919

**Authors:** Harte Singer, Stephen Russell, Andrew Wilson, Mandie Quark, April Coulon, Andrew MacPherson, Kevin Young, Toni Doucette, Scott Ostuni, Maria Marlin, Elizabeth Dods, Matthew Gilbert Koons, Joshua Birkebak

## Abstract

Mushroom forays present unparalleled opportunities to record macrofungal biodiversity in a place and time. The 2024 NAMA annual foray at the Cispus Center near Randle, Washington was the first mass barcoding activity performed by the NAMA DNA Sequencing Committee (DNAMA). The workflows and methods utilized are presented along with planned improvements. Ultimately, 803 ITS barcodes were generated (83.7% of those attempted) for 513 unique taxa belonging to 209 Genera. Of these, 18 novel putative species-level operational taxonomic units (OTUs) were discovered as well as 28 new distribution reports at state level or above. Methods and suggestions for replicating and improving mass barcoding at forays are discussed.

## INTRODUCTION

The North American Mycological Association (NAMA) has been holding annual forays going back to 1961 (excepting 2020 during the COVID pandemic). Originally, only species lists were kept with no organized structure for vouchering or confirmatory identification. In the late 1990s (1997; Copper Mountain, CO) the Voucher Collection Project (VCP; organized by the Voucher Program Committee) started photographically documenting and preserving representative specimens for each identified species (and some additional specimens of particular interest). Specimens, along with relevant photographs and metadata, have been deposited at the Field Museum and are searchable (https://collections-botany.fieldmuseum.org/list?f%5B0%5D=sm_CatProject%3A%22NAMA%20Vouchers%22). Beginning in 2014 (Eatonville, WA), some ITS barcoding activities were initiated by one of the authors (SDR). Over the next decade, DNA barcoding activities increased as methods to expedite throughput while drastically decreasing cost (compared to Sanger-sequencing) using the third generation sequencing platform developed by Oxford Nanopore Technologies were implemented (Russell 2023). Beginning with the 2024 foray (Randle, WA) NAMA invested in modernizing and organizing these DNA sequencing activities through the formation of the DNA Sequencing Committee (DNAMA). This committee was tasked with gathering the relevant stakeholders to coordinate activities and integrate barcoding results with the VCP as well as the broader mycological community.

These NAMA annual foray events involve a large number of attendees collecting mushrooms in a relatively small geographic area, either on their own, or with a foray leader who guides a group to an area to look for mushrooms. Mushrooms are then brought to a central location where they are identified and displayed on tables for attendees to view. Traditionally, identification is done by a small team of experts who check species off a list as they come in from foray participants. These species lists often reflect both the level of expertise and specialization of the members of the identification team. In some cases, specimens of select collections are preserved through dehydration and saved for future study. While this method has been relatively successful in measuring fungal biodiversity, it is fundamentally limited by the expertise and experience of those working the ID table. There are many cryptic species of macrofungi that can only be identified with careful microscopic analysis or using DNA barcoding and these are very likely to be overlooked with a simple visual identification. Furthermore, the expertise bias may artificially inflate the diversity of some groups relative to others. By incorporating a rapid, simple, and inexpensive method of tissue sampling followed by DNA barcoding shortly thereafter, it is possible to greatly reduce identification bias and provide meaningful results in a reasonable time frame.

## MATERIALS AND METHODS

### Specimen Collection

Prior to collection, attendees were instructed to document and photograph their fungal specimens in situ using a standardized field slip containing fields for relevant metadata (Figure 1). Additionally, attendees were asked to upload the photographs to the iNaturalist platform (including at least one with the field label) and to include the foray slip identifier verbatim in the observation description field. When possible, attendees were asked to add an iNat observation number to the slip or, if not uploaded contemporaneously, to add their username and approximate time photographed to be searched later. After returning to the annual foray site, collectors deposited their specimens in a “drop-off” staging area in trays or on tables organized by excursion location along with the associated field slips. The Mycota Lab team samples specimens at this stage of the process before triage, though also intercepts some specimens after triage as described below. Additionally, foray participants were asked to target two genera with many type specimens from the region: *Ramaria* Fr. ex Bonord. (Marr & Stuntz 1973) and *Galerina* Earle (Smith and Singer 1964)

**Figure 1.**
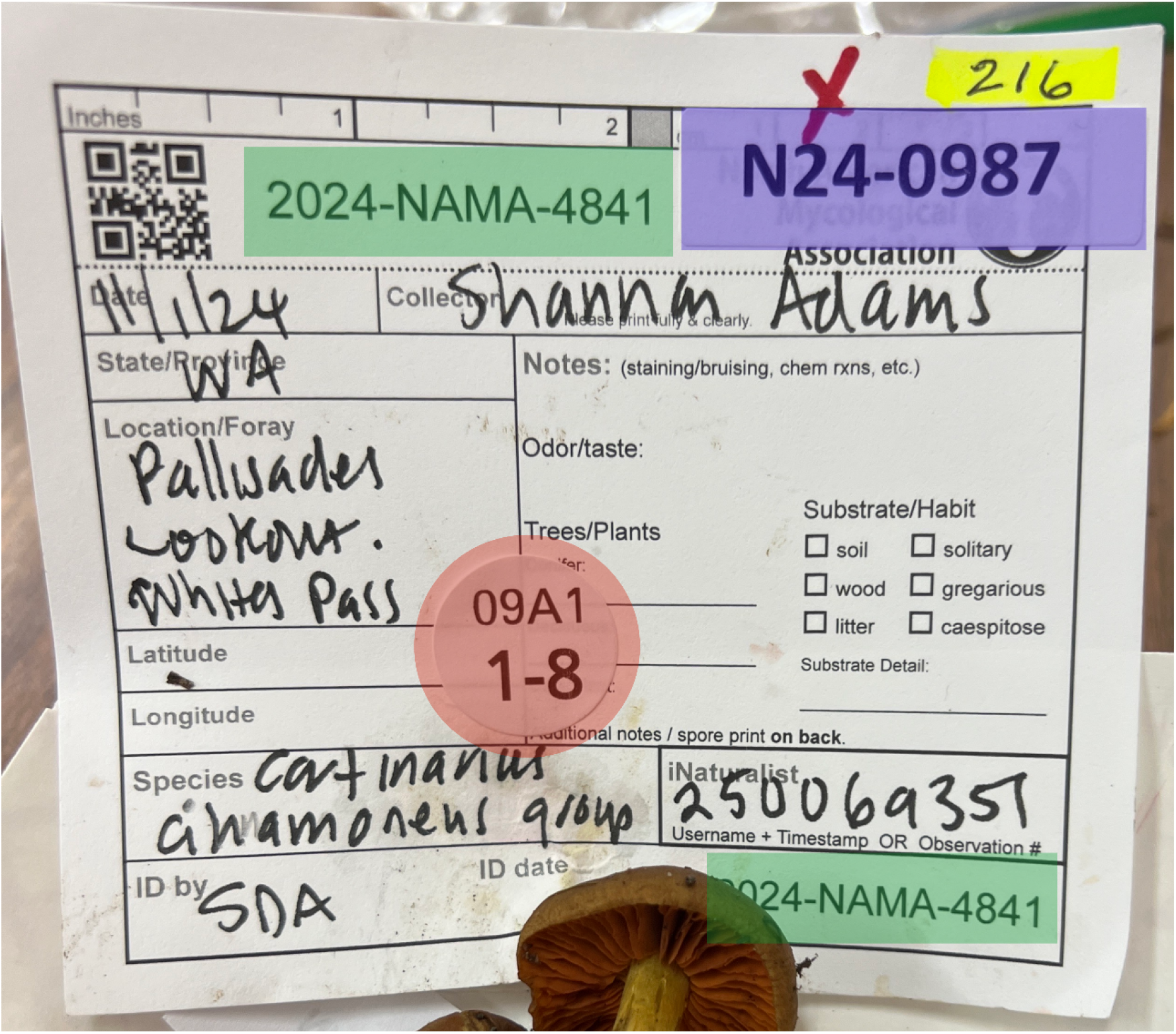
An example of a filled out specimen label. The sample codes and numbers used in the various workflows are highlighted. Green: NAMA Field Slip Number “2024-NAMA-4841”. Red: DNAMA round plate location sticker “09A1 1-8”. Blue: Mycota sample identifier “N24-0987”. Yellow: VCP voucher number “216”.

### Specimen Triage

At the “drop-off” area, specimens were triaged by foray identifiers along with voucher project assistants and other volunteers. Specimens without properly completed field slips (without a location at minimum) or with no field slip at all were discarded. As many specimens as possible were identified to the lowest taxonomic level possible (typically at least to genus). Those identified to species or lower (as well as other specimens of particular interest not identified to species) were screened by the VCP to select a high quality representative specimen for vouchering and preservation at the Field Museum herbarium. Redundant specimens were composited on the display tables or queued for tissue sampling depending on the volume of specimens and rate of processing. Specimens of particular interest flagged by the foray identifiers or VCP team were fast-tracked for tissue sampling and vouchering. All specimens in the voucher collection queue were sampled before accessioning when possible, but many had to be sampled after the fact due to time constraints.

### Tissue Sampling

Volunteers were provided with individually-wrapped unflavored bamboo toothpicks (Blue Top Brand, China), as well as 0.2 uL 8-strip Polymerase Chain Reaction (PCR) tubes with individual swing-caps (Alkali Scientific, Fort Lauderdale FL, USA) filled with a common DNA extraction buffer (100 mM Tris pH 8.0, 19.6 g/L KCl, 10 mM EDTA pH 9; Russell & Singer 2025). For each sample, volunteers were instructed to peel away the cellophane from the toothpick without touching the tip, then dip the tip of the toothpick into the hymenium (fertile surface) of each sporocarp (mushroom) and then dip the toothpick into the buffer and swirl for 2 seconds before discarding the toothpick and closing the tube. An ideal sampling technique would result in a very slight amount of visual tissue on the rough surface of the toothpick, but not a substantial “chunk” of tissue. A corresponding sticker indicating the plate and well position was placed on the NAMA field slip. Each specimen was photographed using the volunteer’s personal smartphone and a unique specimen identification code was marked on a sheet of paper on a line corresponding to the PCR tube in a 96-well plate format. The four digit NAMA field slip number was included in the observation description field.

### DNA Extraction and PCR

After sampling, DNA extraction and PCR was performed on-site on a make-shift lab bench according to the Nanopore protocol used by FUNDIS (Singer and Russell, 2024).

### Data organization

Volunteers were instructed to write the four digit numerical portion of each NAMA voucher number on an array that represented the layout of the PCR tubes in each PCR tube rack. Additionally, they added that same four digit number to the notes field when uploading to iNaturalist. Each tube was assigned the four digit NAMA voucher number in addition to the position in the 96 tube array and array number so that even if the same NAMA number was repeated, the tube-array number would serve as a unique sample identifier. To assign the sequence from each sample tube to an iNaturalist observation, a bulk data export was retrieved using the native iNaturalist exporting tools which included every mushroom observation made within a ∼50 km radius of the Cispus Learning Center over during a 5 day period including the day before the event and the day after the event. Exported metadata included the description (=notes) field as well as the Voucher Number(s) and Voucher Number observation fields. The export (.csv) was sorted to list rows that contained values in either the description, Voucher Number(s), or Voucher Number column and using the “split text to columns” operation, four digit NAMA voucher numbers were isolated. Cells containing these numbers were combined into a single column and only records containing a value in the combined column were used. These were then sorted by the time the observation was made. This filtered and sorted dataset was then used to populate the shared Google Sheet where analysis took place using a Vlookup function to match the sequence record (identified with the four digit NAMA number) to the first observation record that appeared in the iNaturalist data export. In the case where both a field observation and a table observation were made, the first one to be encountered by the Vlookup function would be the field observation.

### Sequence Validation

After the event, barcode validation and identification was performed by the Volunteer Sequence Validation Corps (VSVC) according to the methods of Singer (2024). In brief, the MycoMap (https://mycomap.com) platform was used to apply data quality screening, organize the project data, and BLAST the resulting Nanopore barcode sequence data. Volunteers then individually or collaboratively validated each sequence by comparing the ITS barcode with the source specimen and all other DNA sequences in the database to screen out errors and identify the specimen to a species or *cryptonomen temporarium* (“temp. code”) whenever possible. While doing this, table observations were marked as “casual” by flagging the location as inaccurate on the observation. Before publishing the sequence to iNaturalist, an attempt to find an associated field observation was made. When present, the sequence data was associated with the field observation over the table observation as much as possible but this could not always be accomplished. Sequences were uploaded to GenBank after all validation criteria had been met.

## RESULTS

### Analysis of Workflows

While DNAMA, VCP, and Mycota share overlapping goals and steps, the workflows diverge as soon as the specimens arrive at the foray tables (diagrammed in Figure 2). Mycologists associated with Mycota Labs often photographed the specimen along with its field label and added their own unique sample code. Tissue samples (barcode-quality specimens) were often sampled before collection triage by foray identifiers and the voucher team. Mycota Labs dried and bagged all samples which were shipped off-site to be processed for DNA extraction and sequencing at a later date. After the foray, observations were made based on photographs from the foray table (a.k.a., “table observation”) and associated with the herbarium specimens and, ultimately, DNA sequences.

**Figure 2.**
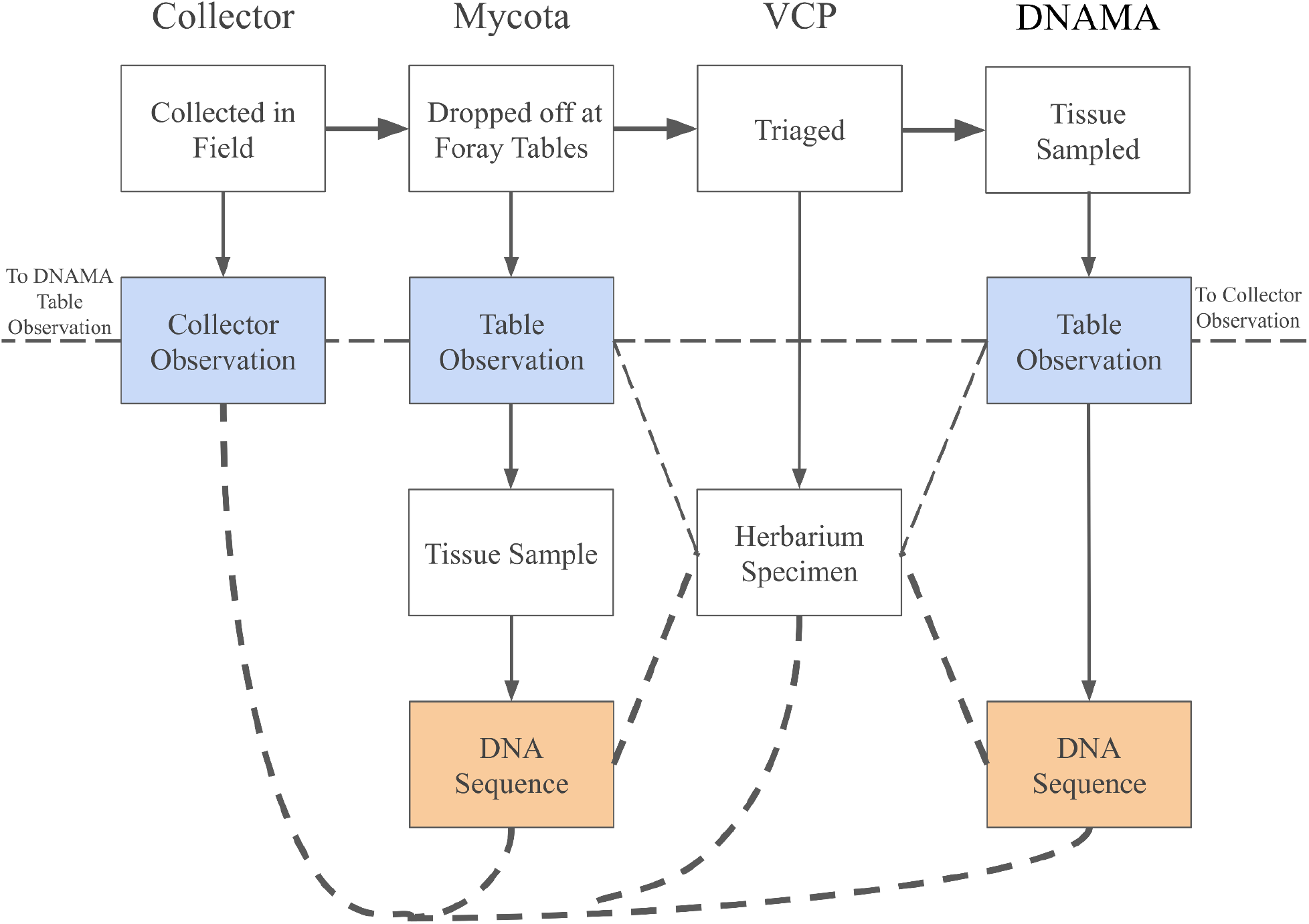
Overview of each teams’ workflows illustrating the journey of a specimen from the field through the sorting process along the top row. Vertical lines represent the direct associations within each workflow. Narrow dashed lines represent manual associations made between workflows after the fact in order to ultimately facilitate the desired associations illustrated by bold dashed lines (i.e., associating the field observation, DNA sequence, and herbarium specimen). Redundancy between workflows are illustrated by shared colors, namely blue for observations and orange for DNA sequences.

In the other workflows, specimens are first triaged by foray identifiers who assign a taxonomic identification to the lowest rank possible. One specimen of each taxon identified to species or lower is vouchered as a museum-quality specimen by the VCP. The careful attention to detail in the documentation required for a museum-quality herbarium specimen limits the capacity to preserve all of the specimens collected, though many specimens identified to genus or higher are vouchered, if of particular interest.

While sampling was somewhat haphazard, the DNAMA team had a specific goal to sequence as many VCP herbarium specimens as possible. Non-herbarium specimens were usually sampled after initial triage by identifiers so that particularly interesting finds could be fast-tracked to the sequencing queue. While using a toothpick to sample fresh material from each specimen into a well on a 96-well plate prefilled with X-amp solution, a table observation was made with a four digit code in the description. This code links the observation with the position on the well plate but no specimens were saved. Plates of tissue samples were extracted and amplified by PCR during the foray, and then purified and prepared for sequencing on the Oxford Nanopore Technologies MinION sequencing platform using a Flongle flow cell (Singer & Russell 2024). Sequencing was completed *in motu* during the drive from Washington to California. The pooled library was sequenced again using an additional Flongle flow cell in California to ensure adequate coverage.

When the sequences were validated by the VSVC, the best attempt was made to use the 4-digit code NAMA field slip code, Mycota sample code, and VCP voucher number to associate the table observation with the collector’s observation, Mycota Lab table observation, and VCP voucher specimen whenever possible. This association was done manually, after the fact and involved members of each team linking 2–3+ observations of the same specimen. The ultimate goal of the teams was to associate the field observation, herbarium specimen, and ITS barcode sequence with each other.

### Sequencing Results

A total of 959 sequencing reactions were performed on specimens from the NAMA foray (Figure 3). Overall, 803 (83.7%) sequencing reactions yielded validated ITS sequences matching the specimen sampled and belonged to 209 genera across 101 families in 27 orders (Figure 4). Of these ITS sequences, 533 were identified to a validly published species name while 268 were identified to a temp. code. These sequences belong to 513 taxa of which 18 were previously unregistered temp. codes. One new continent and one new country distribution record were found as well as seven distribution records for the Pacific Northwest region and 19 new records for the state of Washington. *Cortinarius* (Pers.) Gray and *Inocybe* (Fr.) Fr. were the two most barcoded genera with 53 specimens of each but the former genus had more species level diversity than the latter (Figure 5).

**Figure 3.**
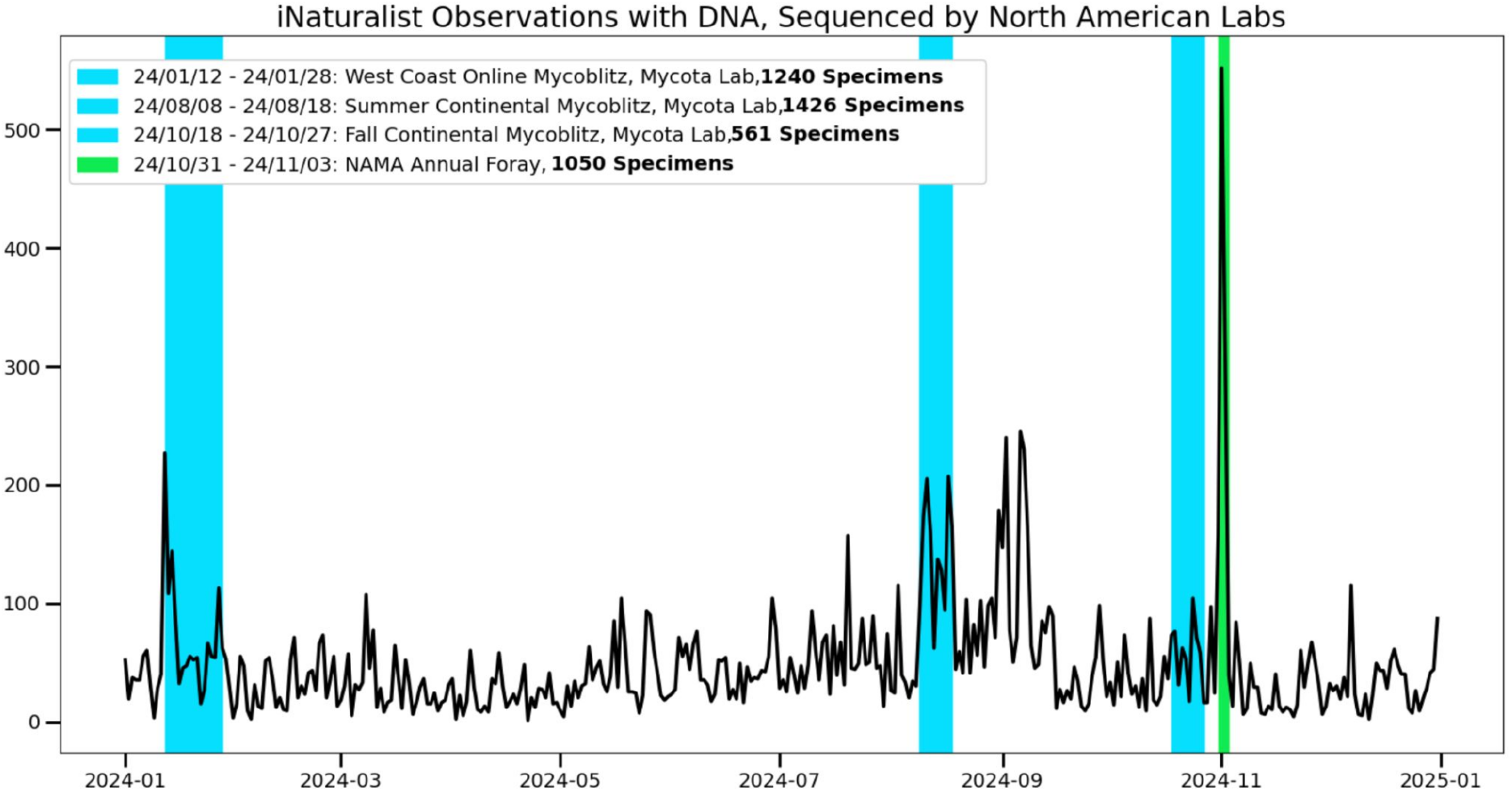
Number of observations made in 2024 with ITS barcode fields on the iNaturalist platform by day of collection over the year 2024 (data accessed 4/2025). The 2024 NAMA foray is shown in green compared to Mycota Lab Mycoblitzes. Inset: Total number of observations with ITS barcodes collected over the length of the event.

**Figure 4.**
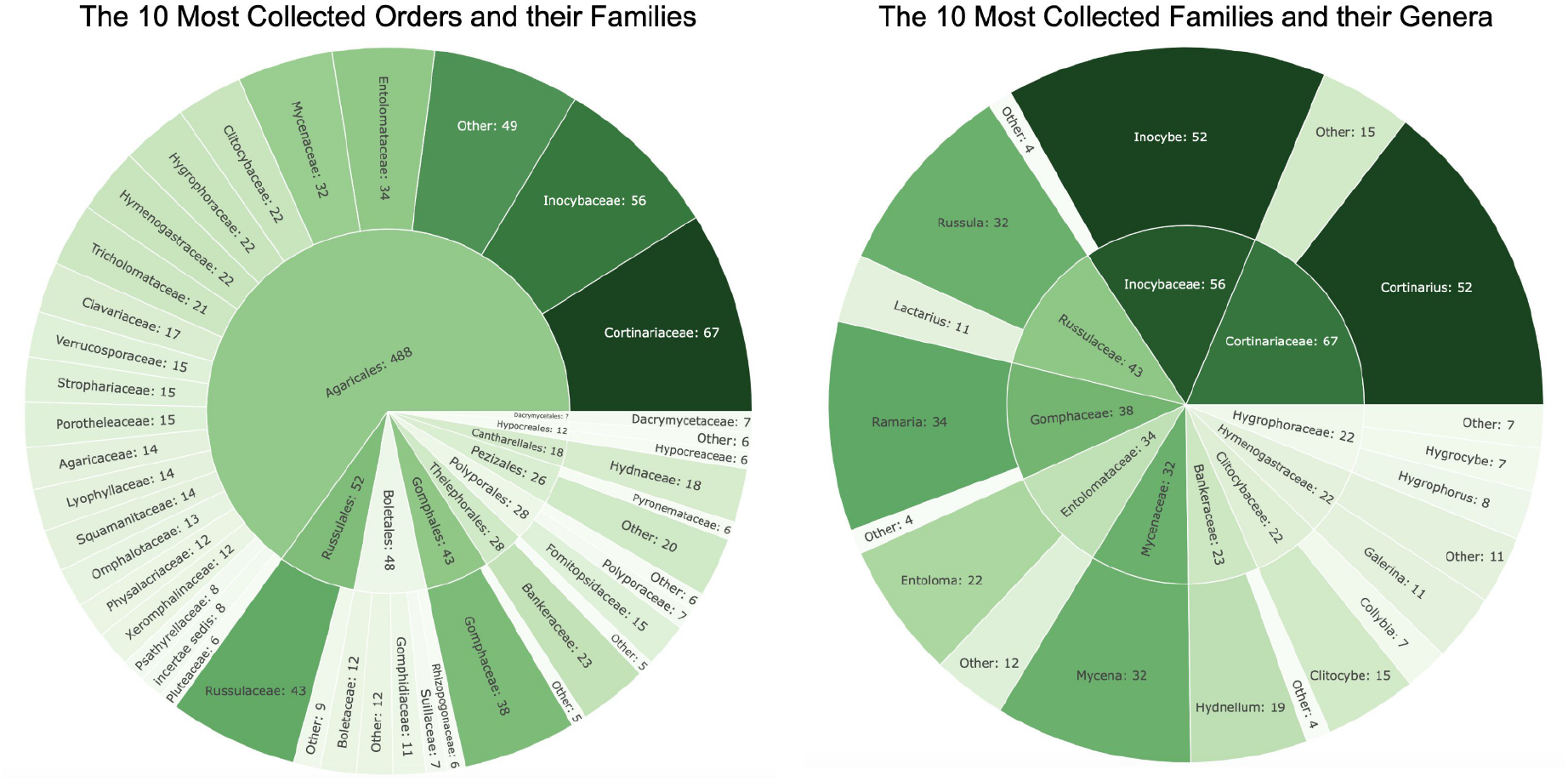
Top ten most Orders (left) and Families (right) with the most sequenced specimens broken down into the next lowest taxonomic rank.

**Figure 5.**
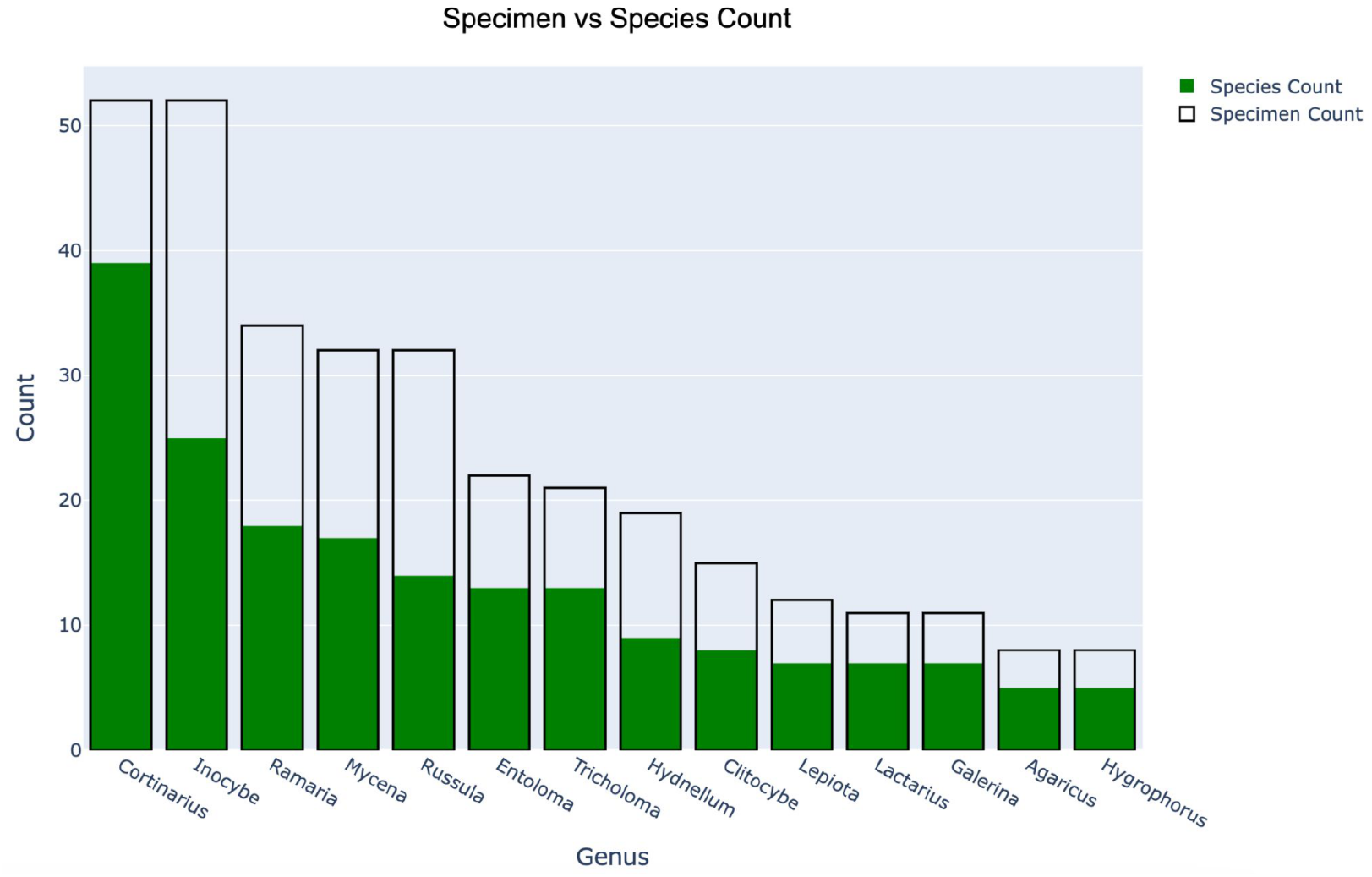
The count of barcoded specimens by genus of the top 14 genera (outlined bar) as well as the number of individual species per genus (green, filled in portion of the bar).

## DISCUSSION

### Some Noteworthy Finds

Of the 801 sequences generated, 18 represented novel OTUs with no matches in our collection of fungal databases (NCBI, UNITE v7.1, iNaturalist, and Mushroom Observer) at the time of analysis (Figure 6). Two of these records represent taxa that are likely to have been described based on commentary from experts on iNaturalist, *Melampsora occidentalis* H.S. Jacks., a rust pathogen on cottonwood leaves, and *Chloroscypha seaveri* Rehm ex Seaver, a minute, olivaceous cup fungus on Western Redwood bracelets (Figure 6A). The remaining 16 represent taxa that have not yet been DNA barcoded and are potentially undescribed, yielding an estimated 2% recovery of novel OTUs from a relatively small geographic area during a restricted time frame. Only one of the 34 *Ramaria* specimens barcoded had no matches, a bright pinkish orange species closely related to *Ramaria longispora* Marr & D.E. Stuntz (Figure 6B). A beautiful silver-purple *Hypoxylon* Bull. matched only a collection made previously from California but was otherwise distantly related to anything else (closest pairwise similarity of 92%; Figure 6C). A small, orange species in the *Pholiotina brunnea* (J.E. Lange & Kühner ex Watling) Singer group, probably had the usual velar remnants washed off by the incredibly wet weather (Figure 6D). *Flammulaster* Earle appears to be a woefully understudied and difficult genus in the family *Tubariaceae*. It is not a huge surprise then that this minute, humicolous specimen (iNat 250635844) turned out to be undocumented, although it may be closely related to *Flammulaster carpophilus* var. *subincarnatus* (Joss. & Kühner) Vellinga (Figure 6E). A small, off-white *Mucronella* Fr. collected from the Cispus center was less than 90% similar to the closest match in our collection of databases., though it should be noted that large differences between species in this genus are common (Figure 6F). Such *Mucronella* specimens are typically referred to as *Mucronella calva* (Alb. & Schwein.) Fr., but there is a huge diversity going under this name across the world.

**Figure 6.**
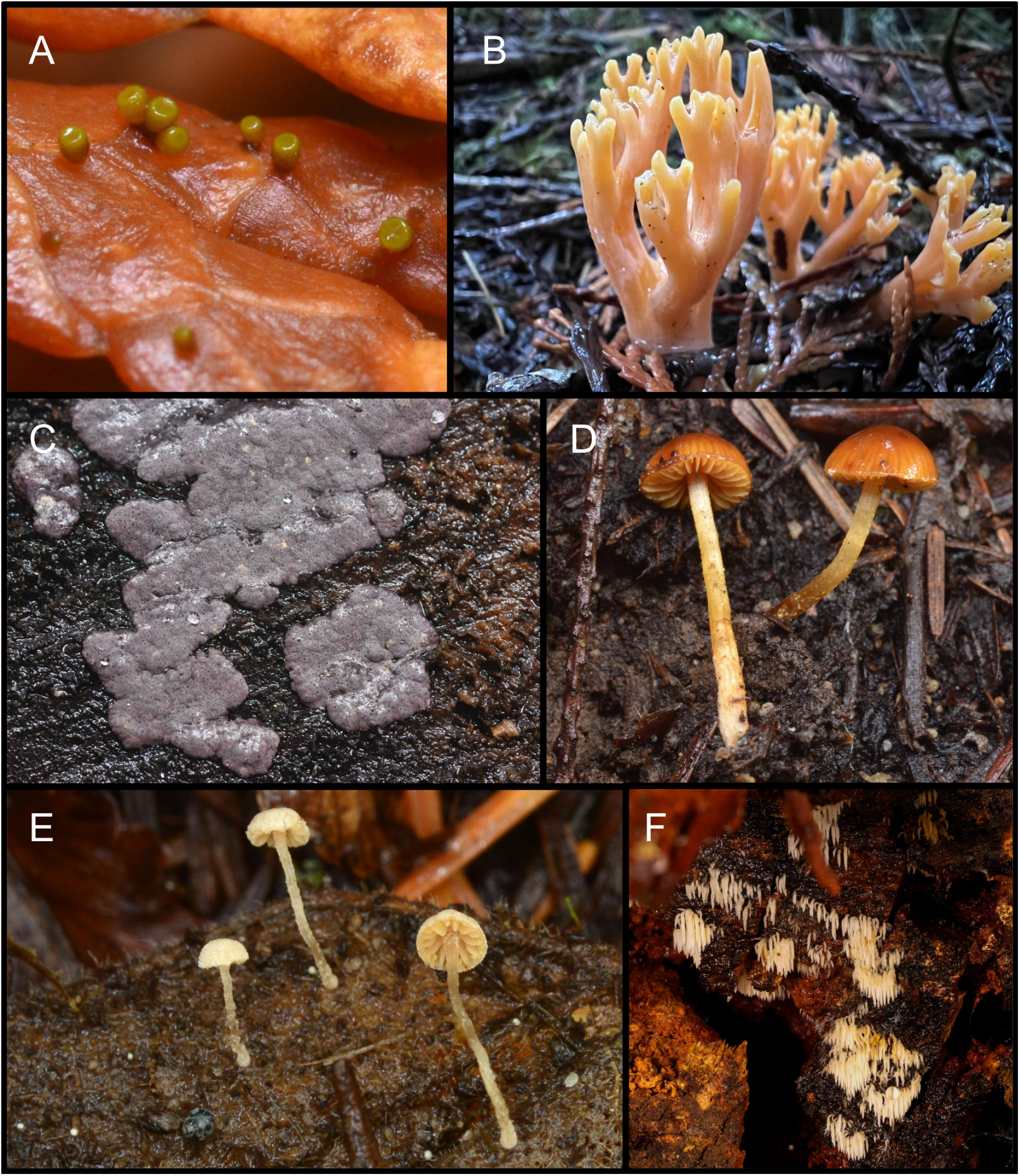
Select specimens for which there were no matching ITS sequences in our collection of public databases, NCBI, UNITE v7.1, iNaturalist, and Mushroom Observer.. Photo credits can be found in the acknowledgements. A. *Chloroscypha seaveri* - iNat 251227326, B. *Ramaria* “sp-WA02” - iNat 250477429, C. *Hypoxylon* sp. ‘WA01’ - iNat 250649956, D. *Pholiotina* sp. ‘WA01’ - iNat 250635842, E. *Flammulaster* sp. ‘WA01’ - iNat 250635844, F. *Mucronella* sp. ‘WA01’ - iNat 250213407.

Additionally, 28 of the sequences were representative of taxa that had not been previously molecularly confirmed from Washington state (Figure 7). Two new distribution confirmations (and one new temporary code) were found during Heather Dawson’s canine assisted truffle foray: *Hymenogaster parksii* Zeller & C.W. Dodge described from California (Figure 7A) and *Melanogaster natsii* Y. Wang, K. Tao & B. Liu described from Oregon (Figure 7E). A fascinating mycophagous fungus, *Entoloma byssisedum* var. *microsporum* Esteve-Rav. & Noordel., may be more commonly encountered than documented as its hyphal stage appears to grow on a diversity of mushrooms (like on the *Russula* Pers. pictured in Figure 7B) and may be dismissed as a species of *Hypomyces* Fr. A unique plant pathogen on Salal (*Gaultheria shallon*) leaves forming tiny white cups, *Lachnum gaultheriae* (Ellis & Everh.) Zeller, was sequenced for the first time ever from the foray (Figure 7C and 7D). A striking coral pink pathogen of turf grass, *Laetisaria fusiformis* (Berk.) Burds., was originally described from Australia but may be widely introduced by humans across the globe (Figure 7F); This specimen (iNat 250080890) represents the first molecular confirmation of the species from North America. A lovely little *Clavulinopsis* Overeem in the *Clavulinopsis corniculata* (Schaeff.) Corner group had previously only been known from the Northeastern United States (Figure 7G).

**Figure 7.**
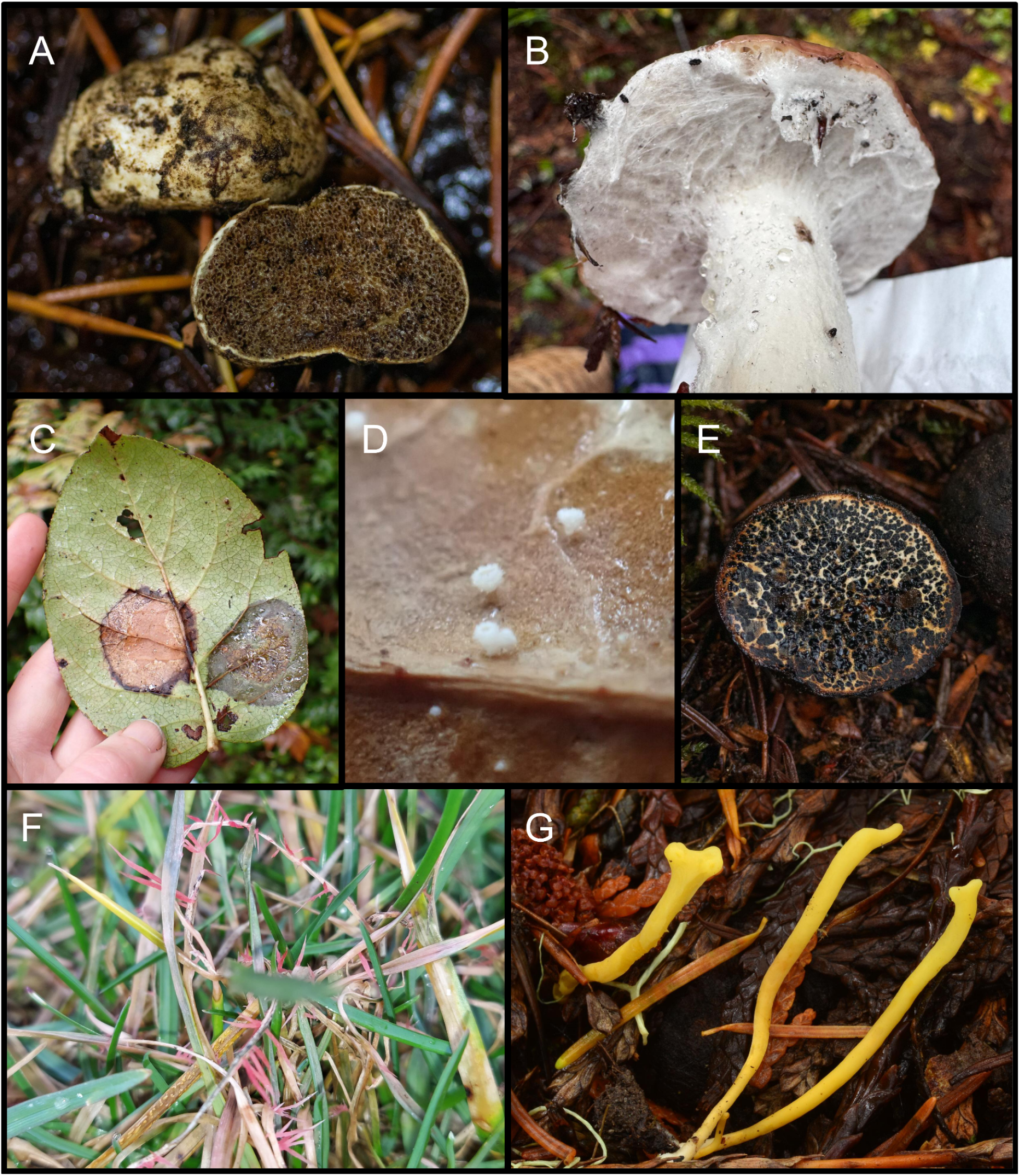
Select specimens that represent new molecularly confirmed distribution reports. Photo credits can be found in the acknowledgements. A. *Hymenogaster parksii* - iNat 250229241, B. *Entoloma byssisedum* var. *microsporum* - iNat 250069566, C. *Lachnum gaultheriae* - iNat 250232440, D. Previous specimen but viewed through a stereo microscope. E. *Melanogaster natsii* - iNat 250098499, F. *Laetisaria fuciformis* - iNat 250080890, G. *Clavulinopsis* “corniculata-NY01” - iNat 250649651.

### Sampling Strategies and Collection Bias

From eleven collections of *Galerina*, eight unique species were identified, three of which were assigned temp. codes based on ITS matches to extant collections. No new temp. codes were created.. In the genus *Ramaria*, 34 specimens were sequenced representing 18 taxa (including temp. codes), one of which had no match to an existing name or temporary code.

The majority of novel taxa were collected by experienced mycologists suggesting a significant sampling bias associated with the recovery of novel data. In comparison, SOMA Camp — a mushroom foray in Sonoma County, California — had a new OTU discovery rate of approximately 3.5%. In both cases, most of the novel OTUs were from collections made by experienced mycologists, which suggests that sampling bias plays a strong role in discovery of new taxa. Additionally, the majority of novel OTUs and new distribution reports were attributed to diminutive, singleton, hypogeous, or otherwise obscure collections, such as the *Gautieria* sp. found using trained dogs. We intend to iteratively improve our sampling workflows to capture as much diversity as possible. Several targeted sampling recommendations for both collectors and DNA triagers are being considered including focusing on specimens that do not get confidently identified, targeting things that grow solitary (and thus less likely to be formally vouchered), and encouraging participants to sample small and obscure fungi.

### Workflow Improvements

An immediate issue identified by the DNAMA team was the amount of time and effort required to upload observations at the time of sampling at the foray. While the Mycota Lab team generated many more dried samples, the time consuming process of tissue sampling had to be done later, off site. The sequencing coordinator (HS) was able to run the Nanopore sequencing reaction on the drive home from the foray, greatly reducing the amount of time spent organizing and sampling specimens. While the VCP process is meticulous and generates data-rich, museum-quality specimens, it is the slowest of the three workflows. Neither teams’ workflow directly interfaced with the VCP, but an attempt was made to sample every vouchered specimen.

In some cases, high-value specimens were sampled by Mycota prior to entering the VCP pipeline which resulted in vouchers that were missing considerable tissue. In other cases there were multiple iNaturalist observations of the same specimens from the ID table. This process could be streamlined by integrating toothpick sampling, tissue harvesting, and photography. Strong communication with the VCP at the point of specimen triage would help prevent excessive damage to potential museum collections.

Whenever possible, table observations were linked to data-rich field observations. The fact that each of the three workflows used different identification codes led to complications manually associating observations with one another. Additionally, reliance on manual data entry could be much improved by image processing and linkage using a QR code on the NAMA field label filled out by each collector. One particular shortcoming of our workflow was our failure to record the location and habitat information from specimens during the sampling or validation process. Automatic image processing solutions would be necessary to bulk process the large number of specimens planned for future events.

Another workflow improvement often asked for by foray organizers and attendees is easily accessing the data generated by barcoding activities. All of the samples for which DNA barcoding was attempted can be found in a iNaturalist project (include both those sampled by DNAMA and Mycota Labs) though it is not as easily organized or sortable in ways that could be desired. In future years, a dashboard could be created to visualize and organize the data according to fields that might be of most interest to foray attendees (e.g., foray number/location, taxonomic group, collector). In the meantime, observations with their associated data can be found here: https://www.inaturalist.org/observations?project_id=220216.

## Glossary

Operational taxonomic unit (OTU): A putative species-level taxon defined by molecular barcode distance.
Cryptonomen temporarium (Temp. code): A functional code name applied to species-level taxa that cannot be positively assigned a published name based on a combination of molecular evidence and expert judgement taxing morphology, ecology, geography into account.
Table observation: An *observation* made *ex situ* on a foray ID table, not in its natural habitat.

## ACKNOWLEDGEMENTS

This was all made possible through the hard work of the NAMA Foray Planning Committee led by Ken Buegeleisen. DNA barcoding activities at the annual foray were funded by NAMA and made possible through thanks to the generous foray registrants who volunteered their time. Of course, the most important contributors were all of the collectors (or at least the ones that filled out their field slips correctly).

Thanks to the following collectors whose Creative Commons licensed photographs were used in this report: Connor Dooley (“corndog”) - 6A, 6C, 6D, 6E, and 7G; “annamycete” - 6B; Matthew Koons (“mgkoons”) - 6F and 7F; Heather Dawson (“heatherdawson”) - 7A and 7E; Krista Willmorth (“fungalforager”) - 7B; and Mariah Rogers (“mkremedios”) - 7C and 7D.

## Notes

https://www.inaturalist.org/projects/nama-2024-dna-barcoding

https://www.inaturalist.org/projects/dnama-volunteer-sequence-validation-corps

### Competing Interest Statement

The authors have declared no competing interest.

